# The importance of geometry in the corneal micropocket angiogenesis assay

**DOI:** 10.1101/193433

**Authors:** James A. Grogan, Anthony J. Connor, Joe M. Pitt-Francis, Philip K. Maini, Helen M. Byrne

## Abstract

The corneal micropocket angiogenesis assay is an experimental protocol for studying vessel network formation, or neovascularization, *in vivo*. The assay is attractive due to the ease with which the developing vessel network can be observed in the same animal over time. Measurements from the assay have been used in combination with mathematical modeling to gain insights into the mechanisms of angiogenesis. While previous modeling studies have adopted planar domains to represent the assay, the hemispherical shape of the cornea and asymmetric positioning of the angiogenic source can be seen to affect vascular patterning in experimental images. As such, we aim to better understand: i) how the geometry of the assay influences vessel network formation and ii) how to relate observations from planar domains to those in the hemispherical cornea. To do so, we develop a three-dimensional, off-lattice mathematical model of neovascularization in the cornea, using a spatially resolved representation of the assay for the first time. Relative to the detailed model, we predict that the adoption of planar geometries has a noticeable impact on vascular patterning, leading to increased vessel ‘merging’, or anastomosis, in particular when circular geometries are adopted. Significant differences in the dynamics of diffusible aniogenesis simulators are also predicted between different domains. In terms of comparing predictions across domains, the ‘distance of the vascular front to the limbus’ metric is found to have low sensitivity to domain choice, while metrics such as densities of tip cells and vessels and ‘vascularized fraction’ are sensitive to domain choice. Given the widespread adoption and attractive simplicity of planar tissue domains, both *in silico* and *in vitro*, the differences identified in the present study should prove useful in relating the results of previous and future theoretical studies of neovascularization to *in vivo* observations in the cornea.

**Author summary:** Neovascularization, or the formation of new blood vessels, is an important process in development, wound healing and cancer. The corneal micropocket assay is used to better understand the process and, in the case of cancer, how it can be controlled with drug therapies for improved patient outcomes. In the assay, the hemispherical shape of the cornea can influence the way the vessel network forms. This makes it difficult to directly compare results from experiments with the predictions of mathematical models or cell culture experiments, which are typically performed on flat substrates or planar matrices. In this study, we use mathematical modeling to investigate how the hemispherical shape of the cornea affects vessel formation and to identify how sensitive different measurements of neovascularization are to geometry.

## Introduction

Neovascularization, or new blood vessel formation, is an important process in development, wound healing, cancer and other diseases. The corneal micropocket angiogenesis assay, shown in Fig 1, is widely used for studying neovascularization *in vivo* [1–3]. The assay involves the implantation of a pellet containing pro-angiogenic compounds into the cornea of a small rodent and observation of the resulting vessel formation over time. Although the cornea is normally avascular, the assay remains popular due to the relative ease with which neovascularization can be observed in the same animal at multiple time-points. In rodents, the cornea is hemispherical, with a thickness on the order of 10 to 20 vessel diameters [4]. The pellet is not usually located on the pole of the cornea; it is shifted slightly toward the base or limbus. It is evident from experimental images that the geometrical configuration of the cornea-pellet system influences neovascularization patterns [1]. As shown schematically in Fig 1C, new vessel formation is often focused in the region where the distance between the pellet and limbus is smallest, with vessels tending to grow toward the pellet.

**Fig. 1.**
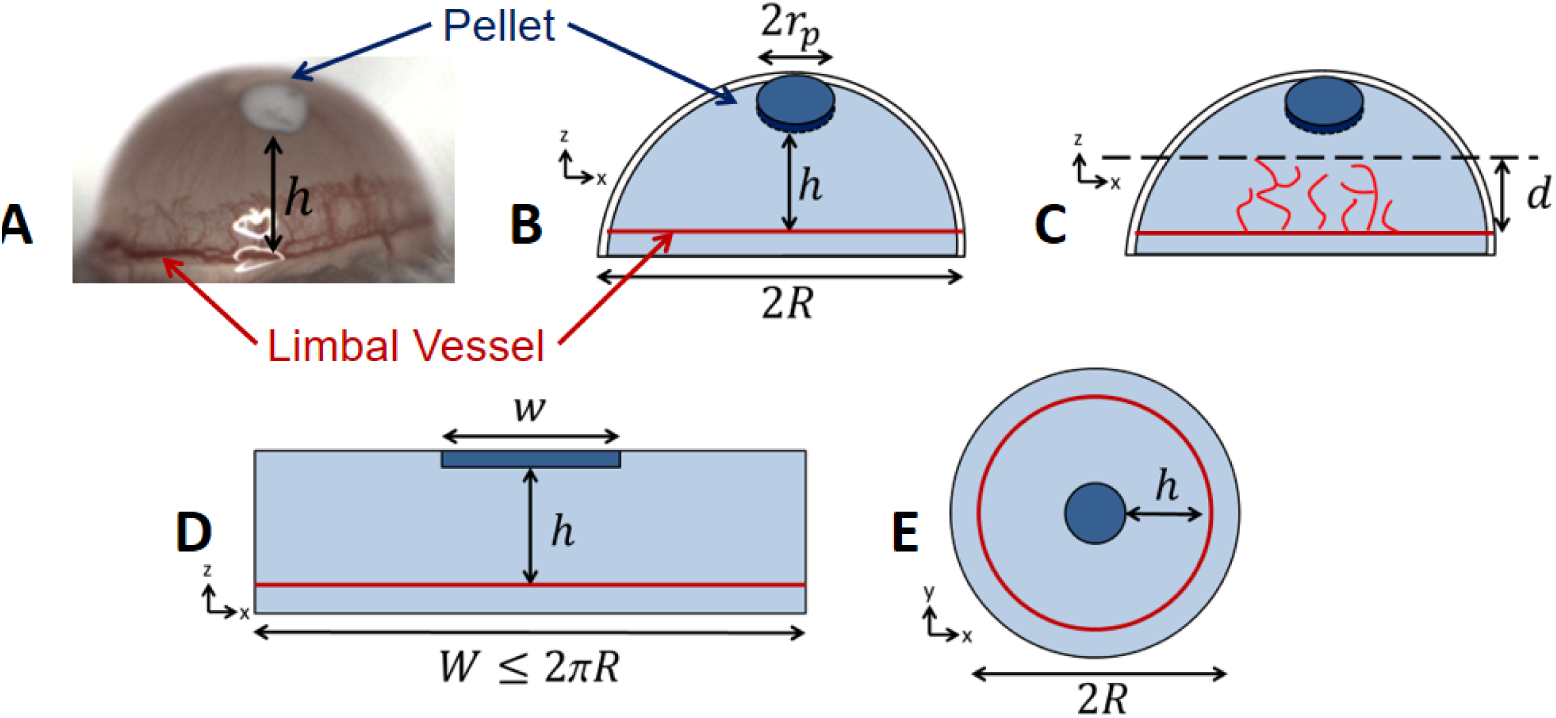
Schematics of the corneal micropocket assay and typical *in silico* and *in vitro* geometries. **A)** An image of the micropocket assay from Connor et al. [5]. **B)** The placement of a pellet of radius *r*_*p*_ into the cornea at height *h* from the limbus, where *h* is the distance along the shown *y* or *z* axis. **C)** The resulting neovascularization, quantified by distance *d* of the front to the limbus. **D)** A typical planar model of the cornea-pellet system, with the pellet width *w* set equal to cornea width *W* for one-dimensional mathematical models. **E)** A typical circular geometry.

The simplicity of the micropocket assay has made it an attractive candidate for comparing vessel network formation with mathematical (*in silico*) and *in vitro* models. As reviewed in Jackson and Zheng [6], many mathematical models of neovascularization have been motivated by the micropocket assay, providing valuable insights into the process [5, 7–14]. To date, these models have exclusively adopted either one-dimensional (1D) or two-dimensional (2D) representations of the cornea-pellet system, with the former allowing efficient, continuum modeling of the developing vessel network by the solution of systems of partial differential equations (PDEs) [5,10, 11].

While most modeling studies are based on qualitative analyses of the assay, some have performed more direct, and even quantitative, comparisons with experimental observations. For example, in a series of studies, Tong and Yuan [3,7,13] developed a model of the assay using a 2D circular domain, as shown in Figs 1 E, based on earlier discrete modeling approaches by Stokes and Lauffenburger [9]. The authors compared predicted patterns of vascularization with their own experimental observations, using a range of metrics such as vessel length, migration distance and projected width of the vascularized region. The authors used their theoretical model to better understand the interplay between diffusible growth factors, growth factor binding to endothelial cells and endothelial cell density, based on observations of vascularization as pellet loading was increased. Harrington *et al.* [14] used a similar modeling approach to study inhibitor loading and positioning in the cornea, with qualitative comparisons of vascular patterning with experiment. Jackson and Zheng [6] developed a detailed, discrete, model of endothelial cell proliferation and migration in a 2D circular domain. The authors performed qualitative and quantitative comparisons of vascular patterning with experimental results from Sholley *et al.* [2]. More recently, Vilanova *et al.* [15] developed a phase-field model of individual vessels and simulated the assay in a 2D circular domain as an element of a more detailed study. The authors performed qualitative comparisons of vascular patterning and front velocity with the previous studies of Tong and Yuan [3,7]. In terms of continuum models, Connor *et al.* [5] used a classical 1D modeling approach to perform detailed quantitative comparisons of predicted vessel densities with their own experimental measurements.

Given: i) the widespread use of 1D and 2D models of the assay, ii) the use of both qualitative and quantitative comparisons between predicted patterning and experiment, and iii) the observation that the geometrical configuration of the cornea-pellet system influences neovascularization patterns in experimental images, it is important to understand how the three-dimensional (3D) geometry of the assay affects vessel network formation relative to the planar tissue domains typically used in mathematical models. While there are many 3D mathematical models of sprouting angiogenesis [16–20], none have focused on the particular geometry of the cornea-pellet system, making it difficult to predict the influence of the assay geometry without a dedicated study. Such a study brings the additional challenges of needing to use a relatively large simulation domain and accounting for the interaction of vessels with curved tissue boundaries at the epithelial and endothelial surfaces of the cornea.

In the present study, we develop a discrete, 3D, off-lattice mathematical model of neovascularization in the cornea-pellet system, focusing on emulating the *in vivo* configuration. We use a simplified treatment of the underlying biology, focusing instead on how the adoption of different geometries, including planar 2D and 3D cultures in rectangular or circular configurations, affects vessel network formation relative to the *in vivo* case. The primary strengths of the study are: i) the simulation of neovascularization in large, 3D tissue domains and ii) the ability to compare predictions across several different geometries and with different biophysical processes activated and de-activated.

As part of this comparison we also identify metrics of neovascularization with high and low sensitivity to the choice of tissue domain. As a result, we can predict which *in vitro* and *in silico* tissue domains most closely resemble the conditions of the *in vivo* experiment and which metrics of neovascularization are most suitable for performing comparisons. Given the widespread adoption and attractive simplicity of planar tissue domains, the differences identified here should prove useful in relating and translating the results of previous and future *in silico* and *in vitro* studies of neovascularization to *in vivo* observations in the cornea.

## Materials and methods

Neovascularization is simulated in seven different tissue domains, representing those typical of *in silico* and *in vitro* modeling studies and the *in vivo* assay. A phenomenological model of sprouting angiogenesis is adopted, motivated by several previous studies [6,7,14,22], but extended to 3D. A ‘soft-contact’ model is also introduced to account for interactions of migrating vessels with the cornea boundaries. It is assumed that the pellet contains a single pro-angiogenic compound, Vascular Endothelial Growth Factor 165 (denoted VEGF in the present study). Two situations are considered, one where the concentration of VEGF in the tissue domain is described by a time-independent, spatially varying field where VEGF levels decrease linearly from the pellet, and the other where VEGF dynamics are explicitly modeled, following approaches in Tong and Yuan [7] and Connor *et al.* [5].

Overviews of the simulated tissue domains, angiogenesis model and VEGF dynamics model are provided in this section.

Simulations are built using the Microvessel Chaste library, which is a collection of C++ classes providing models and numerical tools for creating angiogenesis simulations. The library motivation and high-level design are described in a dedicated publication [21]. Users of the library build their own simulations from the available C++ classes, as has been done in the present study for the particular problem at hand, rather than use it as a monolithic solver. The geometrical solid models, PDE solvers and angiogenesis models described below are all built using the library.

### Tissue domains

Fig 2 shows 3D renderings of each of the studied domains, along with the naming convention used when presenting results. The pellet radius *r*_*p*_ = 200 µm is based on data from Connor *et al.* [5], but reduced from their value of 300 µm to facilitate placement in the cornea. In 3D simulations, pellets are assumed to have a thickness of *T*_*p*_ = 40 µm and are situated mid-way between the epithelial and endothelial sides of the cornea. In 2D simulations, the cornea thickness is neglected, while in 3D a value of *T* = 100 µm is used [4]. The cornea radius is fixed at *R =* 1300 µm, which is a suitable value for mouse [4]. The ‘Hemisphere’ geometry is formed by a 360° revolution of a circular arc of radius *R* and angle 90° about the polar axis, followed by an extrusion through a distance *T* along the inward normal to the revolved surface, giving a 3D volume. The cylindrical pellet is placed inside this volume and is completely enclosed by it. For the Hemisphere, the pellet height *h* above the limbus is the distance as projected into 2D, as would be typically measured in experimental images, rather than the distance along the geodesic from the limbus to the pellet.

**Fig. 2.**
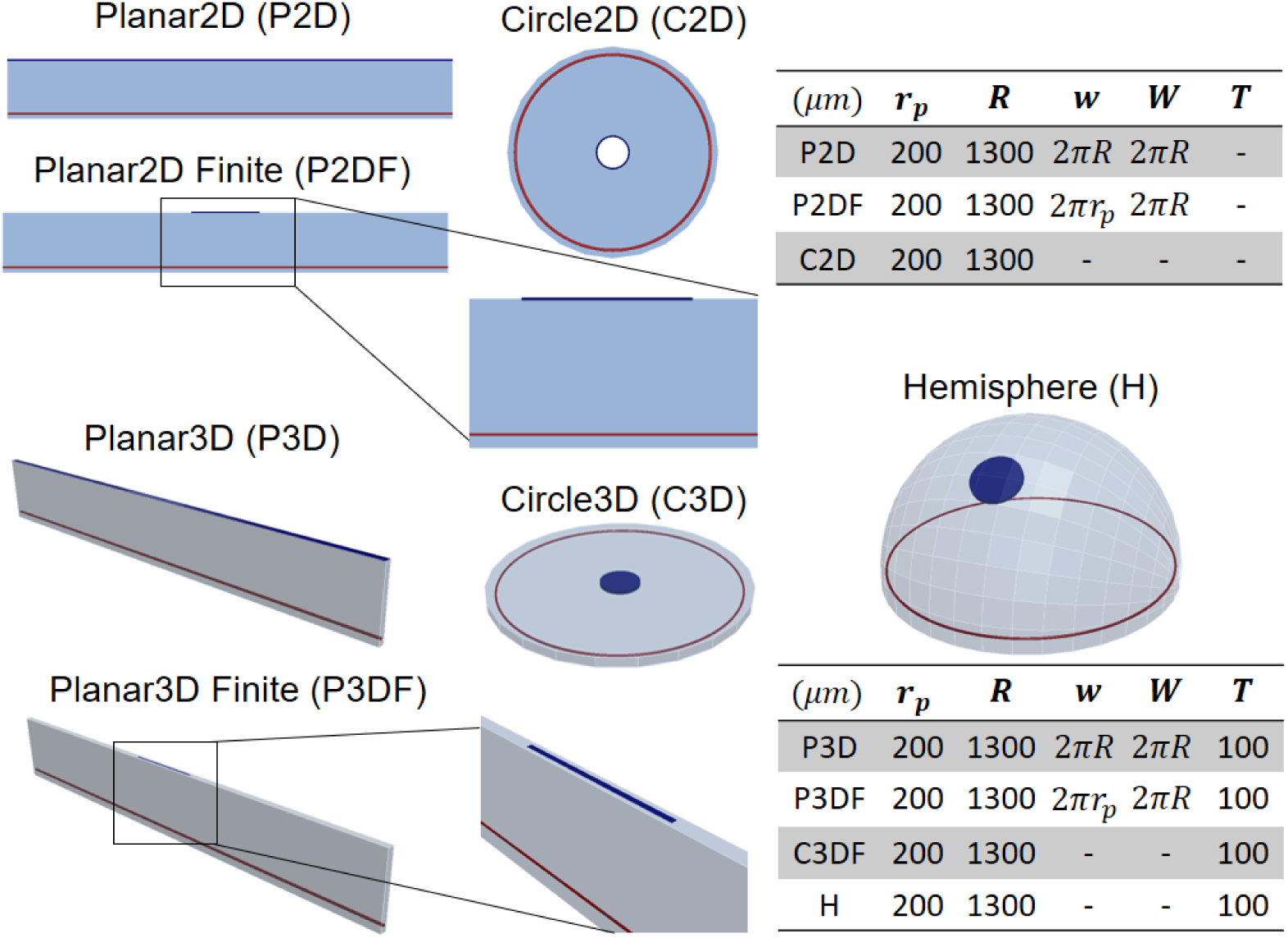
3D renderings of the studied tissue domains, including dimensions. 3D renderings of the studied simulation domains, including adopted abbreviations. Domain dimensions are in µm and follow the notation in Fig 1, and *T* is the cornea thickness.

All simulations begin with a single blood vessel positioned a small height *ε* = 100 µm above the base of the cornea and mid-way between the epithelial and endothelial sides of the cornea. The vessel occupies the entire width (or circumference) of the domain at that position. Vessels are represented as collections of infinitesimally thin, straight-line segments joined at point locations, denoted ‘nodes’, shown schematically in Fig 3. Nodes can be connected to one or more segments. They are assigned numerical or boolean attributes, such as ‘Radius’ and ‘Migrating’ respectively as needed. In the present study line segments can be thought of as corresponding to vessel centrelines. Nodes do not necessarily correspond to individual biological cells, rather a constant number of endothelial cells per unit vessel length 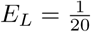 [23] is assumed on each line segment, based on 5 µm radius capillaries.

**Fig. 3.**
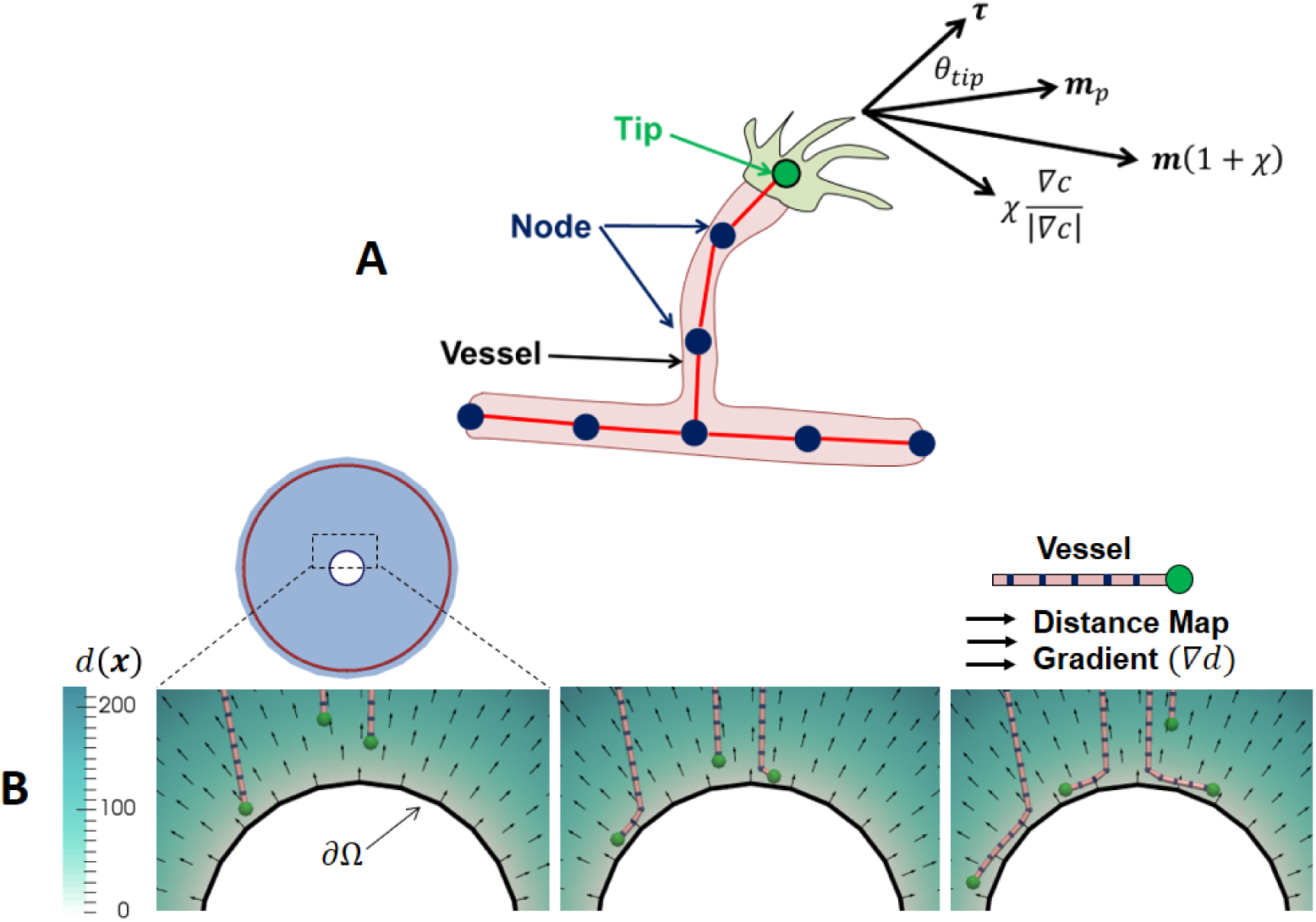
The discretized vessel network representation and an illustrative example of the boundary repulsion model. **A)** The discretized vessel network showing ‘nodes’, ‘vessels’ and ‘tips’. Directions used in the migration rule are also shown. **B)** An illustrative example of the boundary repulsion model, with vessels deflected along a circular boundary. A distance map *d*(*x*) to the boundary (in um) and distance map gradient directions ∇*d* are also shown with arrows. Vessels are only repulsed from the endothelial and epithelial cornea surfaces in the present study, as the pellets are never reached.

### Angiogenesis model

During angiogenesis the vessel network is updated at discrete time intervals Δ*t* = 1.0 h following a sequence of migration, sprouting and anastomosis stages. In a single time step the following stages occur in order: tips migrate, ‘nearby’ tips anastomose, new tips form due to sprouting and any remaining ‘nearby’ tips anastomose.

A simple, phenomenological model of sprouting angiogenesis is used. The average rate of sprout formation at a node located at *x* is [22]:

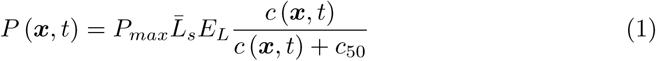

where *P*_*max*_ is the rate of sprouting per cell, *c*(*x*, *t*) is the VEGF concentration at location *x* and time *t*, *c*_50_ is the VEGF concentration at which the rate of sprouting is half-maximal and 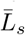 is the averaged length of the two line segments joined to the node. Concentrations at sampled locations are calculated by interpolation from nodal values on finite element meshes using linear triangular or tetrahedral shape functions.

A simple description of lateral inhibition is used, with *P* = 0 within a distance 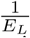 of a node that has already been selected for sprouting. Simulations are discretized in time using a fixed step of Δ*t*. In each time step a random number *z* ∈ [0, 1] is chosen from a uniform distribution at each node and a sprout forms if *z* < *P*Δ*t*. A different random number is generated at each node. Sprouts form in the network by creating a new node at the sprout location and offsetting it by the tip speed *s* times the time increment Δ*t* in a random direction, normal to the parent line segment. A new line segment is created between the new node and the original sprout location and the new node is marked as ‘Migrating’.

Migrating tips (nodes marked as ‘Migrating’), illustrated in Fig 3, are assumed to move at constant speed *s* = 10 μmh^−1^. This speed is chosen so that the average time for the vascular front to reach the pellet is on the order of 4 days, which is consistent with experimental observations [1]. A persistent, off-lattice random walk is used to describe the migration of tips through the extracellular matrix of the stroma [6,7,14]. The migration direction ***m*** is given by:

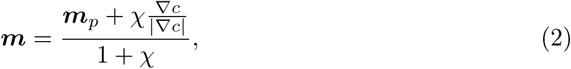

where *χ* is a dimensionless weighting parameter controlling chemotactic sensitivity, ***m***_*p*_ is a unit vector in the persistence direction and ∇*c* is the gradient of the VEGF concentration, calculated using a centered difference approach from nodal solutions on a finite element mesh and sampled at the tip. The random persistence direction ***m***_*p*_ is obtained by rotating the unit tangent vector along the vessel ***τ*** an angle *θ*_*tip*_ away from its original direction in an arbitrary plane, as shown in Fig 3, following similar approaches to account for extracellular matrix interactions in [3,14]. The angle *θ*_*tip*_ is chosen from a normal distribution with zero mean and standard deviation *σ.* The approach for modeling chemotaxis is similarly based on those of previous studies [9,13].

An important distinction from previous studies is that the finite extents of the cornea are accounted for: migrating tips are not permitted to leave the simulation domain. A ‘soft contact’ model is adopted so that tips approaching the boundary of the domain are gradually deflected along the tangent to the bounding surface. Biologically this represents tip cells failing to penetrate the stiffer tissue present on the epithelial and endothelial cornea surfaces, while ‘soft’ contact is chosen for more robust numerics. The strength of the repulsion from the boundary increases as the boundary is approached according to:

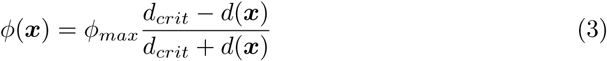

where *d*(*x*) is the minimum distance to the domain boundaries, *d*_*crit*_ is the distance to the boundary at which repulsion is activated and *φ*_*max*_ is the dimensionless maximum repulsion strength. The repulsion biases the motion along the tangent to the bounding surface according to:

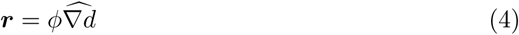

where the 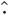 operator produces a random unit tangent. The position of migrating nodes which have not just sprouted is updated from ***x***(*t*) to ***x***(*t* + Δ*t*) at each time step:

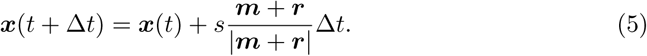

where *s* is the tip velocity and Δ*t* is the time step size in hours. Fixed values of the repulsion strength *φ*_*max*_ = 5 and critical repulsion distance *d*_*crit*_= 25 um are used for all simulations. These values are chosen to ensure a gradual deflection away from the surface, without overly influencing the migration of tips that are far away from the boundaries. An illustrative example application of the boundary repulsion model on a contrived circular domain is shown in Fig 3B.

Endothelial tip cells are known to find and merge with other endothelial tips and immature blood vessels during migration, in a process known as anastomosis [24]. There is still uncertainty about the mechanisms by which they meet, but mechanical and chemical guidance are known to contribute [24]. When moving from 2D to 3D models of sprouting angiogenesis it is necessary to define a region within which tips will merge with vessels and each other to allow for the identification of intersections. In this study, a relatively small radius of *r*_*ana*_ = 5 um is used, which is on the order of the vessel radius. During simulations, sprouting and migration events occur during discrete time intervals Δ*t*. Anastomosis is implemented by identifying the nearest line segment to nodes marked as ‘Migrating’ after each migration or sprouting event. If the distance from the node to the line segment (point to line distance) is less than the radius *r*_*ana*_ an anastomosis event occurs. An anastomosis event can be either a ‘tip-to-tip’ interaction, in which the migrating node is moved to be coincident with its neighbor and both are de-activated, or a ‘tip-to-vessel’ interaction in which the migrating node is moved onto the line segment, de-activated and a new branch is formed. Given the small anastomosis radius used in the present study it is assumed that only biological cells in direct physical contact will anastomose, which may be overly restrictive if mechanical guidance ultimately plays a strong role in the process.

### VEGF dynamics

VEGF dynamics are treated in two different ways in this study. In the first case, the pellet dynamics are ignored and a time independent, spatially varying VEGF concentration field is imposed in the tissue domain. In the second case, the dynamics of VEGF release from a nylon pellet are explicitly modeled. The motivation for the former model is that it allows the effects of cornea geometry on angiogenesis to be observed independently of the pellet representation.

For the first case, a constant VEGF gradient between the limbus and the pellet is applied, given by:

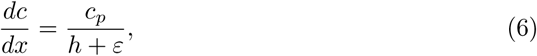

where *x* is a positional coordinate along the geodesic between the limbus and pellet, *ɛ* is a small offset from the base, corresponding to the position of the initial vessel, and *c*_*p*_ is the VEGF concentration in the pellet, which is assumed to be constant in time in this case. By specifying:

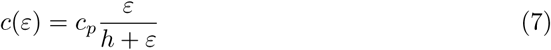

the same concentration and concentration gradient magnitudes are maintained at the limbus in all representations. Aside from the Hemisphere, the concentration at the base is 0 nM and at height *h* + *ε* (the pellet location) it is *c*_*p*_.

For the second case, the transport and decay of the VEGF in the pellet are considered, meaning that *c*_*p*_ can now change over time. The dynamics of VEGF in the pellet are not known in detail, but as per Tong and Yuan [7] a high rate of reversible binding to the nylon pellet constituents is assumed. Under this assumption it is possible to derive the following relationship between the total *c*_*p*_ and free *c*_*f*_ amounts of VEGF in the pellet [5]:

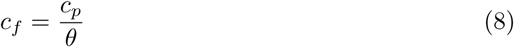

where *θ* ≥ 1 is a dimensionless binding parameter. Free VEGF can decay in the pellet at a rate *λ*_*p*_ or leak through the cornea-pellet interface, which has an effective permeability *κ*_*p*_. It is assumed that the VEGF concentration is spatially uniform within the pellet, which has volume *Ω*_*p*_. Balancing mass leads to the following differential equation describing the time rate of change of VEGF in the pellet:

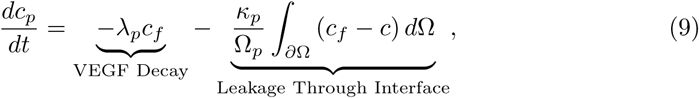

where the integral is over the pellet surface *∂Ω* and c is the concentration of VEGF in the cornea, at the interface. The initial concentration *c*_*p*_(*t* = 0) can be determined from the implanted VEGF mass *m* = 300 ng [5] by:

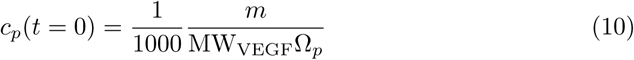

where the VEGF molecular weight MWVEGF is 45 kDa or 45 000 gmol^−1^ and the factor of 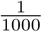 _0_ converts from molm to M.

It is assumed that VEGF diffuses isotropically in the cornea, with a diffusion coefficient *D* = 2.52 x 10-^7^ m^2^h^−1^ [25,26] and decays naturally at a rate *λ* = 0.8 h^−1^ [27]. It is also assumed to enter perfused vessels (and be washed away) and bind to endothelial cells. Combining these processes, we deduce that the dynamics of VEGF in the cornea can be described by the following reaction-diffusion equation:

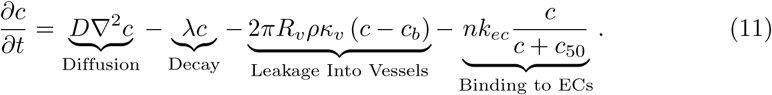

Here *κ*_*v*_ = 3 x 10 mh [28,29] is the permeability of vessels to VEGF,*R*_*v*_ = 5 um [30,31] is the assumed vessel radius, *c*_*b*_ = 0 M is the amount of VEGF in the blood, assuming it is quickly removed, and *ρ* and *n* are respective vessel line and tip densities. The parameter *k*_*ec*_ is the rate of VEGF binding per endothelial cell and *c*_*50*_ is the VEGF concentration at which the rate of binding is half maximal. Continuum reconstructions of the vessel line and tip densities are calculated from the discrete network representation by summing the total vessel length (or number of ‘migrating nodes’) per finite element and dividing by the element volume. These quantities are then used in the calculation of source and sink rates on an element-by-element basis in the finite element solution of the PDEs. Although widely used [6,16], this approach can lead to a PDE and angiogenesis model solution dependence on mesh size. In the present study the ratio of element length to vessel diameter is approximately 3.

The extent to which VEGF will pass through the outer cornea layers is not clear, nor whether it will pass through the epithelial layer and into the aqueous humor or the collagen-rich limbus. It is assumed that the rate of such leakage is low, and no flux boundary conditions are imposed on all outer surfaces of the cornea *∂Ω*_*cornea*_, that is:

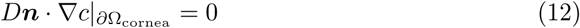

where *n* is the inward surface normal. On the cornea-pellet interface the following mass balance is assumed:

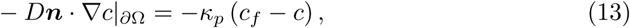

where 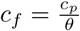 is the amount of free VEGF in the pellet.

Eqns (9)-(13) are solved numerically (detailed below), subject to the initial condition of no VEGF in the cornea. In the 2D geometries, the cornea-pellet interface *∂Ω* is a line of length *w* for the planar case or 2*πr*_*p*_ for the circle. In the Planar3D geometries it is a rectangle of height *T* or *T*_*p*_, depending on whether a finite sized pellet is assumed, and width *w.* In the remaining geometries, the interface is the entire outer surface of the spatially resolved pellet.

### Parameter values

Table 1 summarizes the parameter values adopted in this study. Parameter values with sources denoted as ‘This Study’ are discussed in this section unless previously introduced.

A pellet thickness of *T*_*p*_ = 40 um is used in this study, which is less than the *T*_*p*_ = 60 um value reported in Connor *et al.* [5]. This is to facilitate placement of the pellet in the simulated Hemisphere geometry. The chemotactic sensitivity range *χ* ∈ [0,0.5] is chosen to cover extreme cases where the resulting vessel network is not directed towards the pellet and highly directed towards it. The range of the deviation in persistence angle *σ* ∈ [0, 20] degrees covers cases where straight vessels form, through to cases with vessels with tortuosity similar to that observed in experimental images of the assay. The global time step Δ*t* = 1 h is chosen to give average segment lengths of 10 um, which leads to a physically realistic vessel tortuosity. The initial offset from the limbus of *ε* = 100 μm is in agreement with experimental images [5].

**Table 1.**
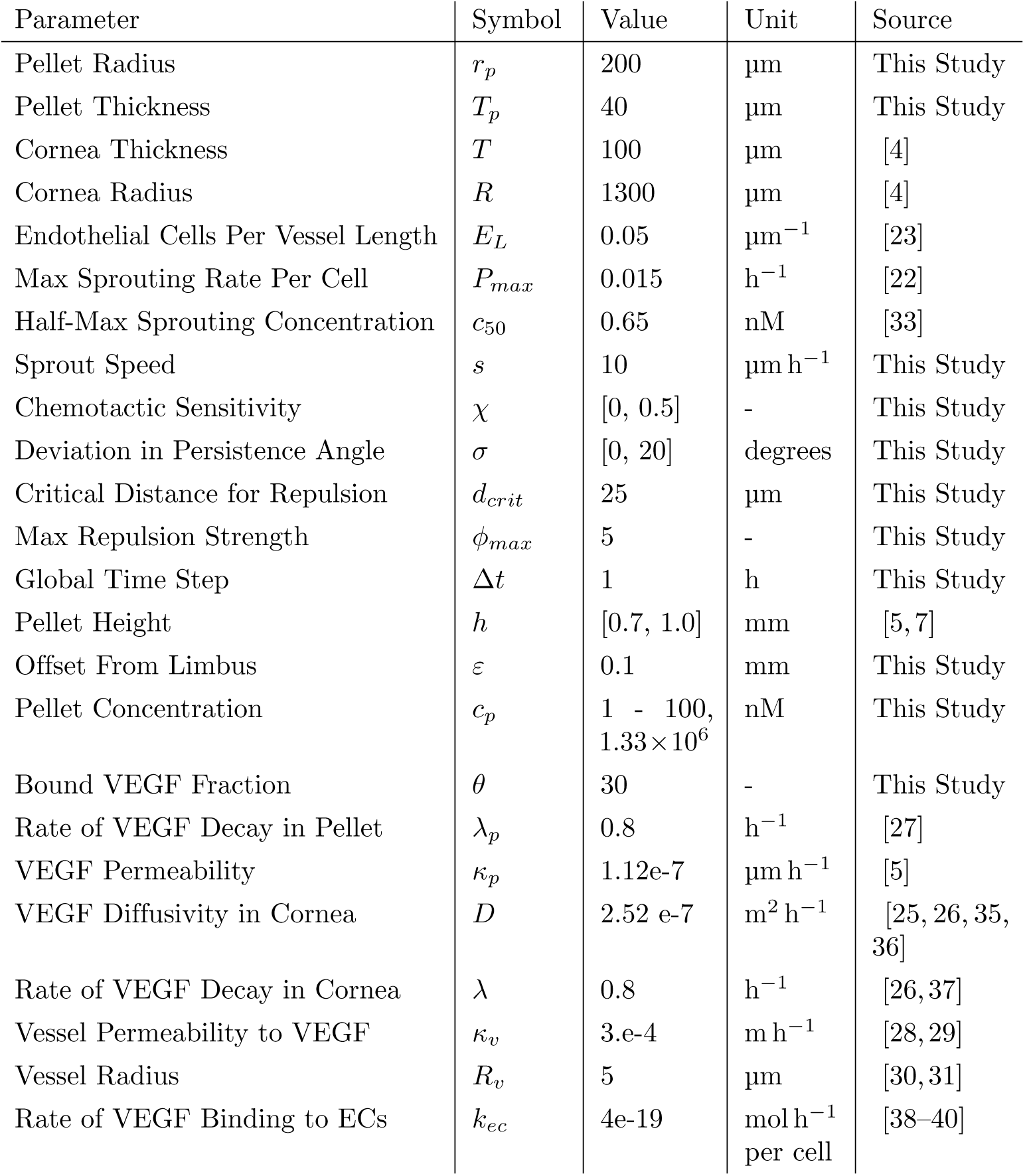
Adopted parameter values for the neovascularization simulations.

The amount of growth factor in implanted pellets is usually known by mass, with a value of 300 ng for VEGF reported in Connor *et al.* [5]. For our pellet volume of 0.0075 mm^3^ and VEGF molecular weight of 45 kDa, this corresponds to a pellet concentration of approximately *c*_*p*_ =1330 μM, which is adopted for the dynamic VEGF model. For the time-independent VEGF model lower pellet concentrations of *c*_*p*_ = 1 to 100 nM are used, which give similar concentrations at the limbus to the dynamic model in the early stages of the simulation. The bound fraction of VEGF in the dynamic model *θ* = 30 is chosen to give a time of VEGF depletion in the pellet of approximately 4 days.

## Simulation details

The VEGF PDE is solved using the finite element method with linear basis functions. A simple forward-Euler time-stepping scheme is adopted, with suitable time steps identified by convergence studies. The maximum PDE solution time step is 0.05 h and a typical grid side length is 30 μm. PDE solutions are updated to the end of the global time step Δ*t* before solutions are sampled for use in the sprouting and migrations rules.

Simulation results are presented in terms of the ‘vessel line density’ *ρ*(***x***,*t*), which is defined as the vessel length per unit volume, and ‘tip density’ *n*(***x***,*t*), which is the number of migrating tips per unit volume. Densities are calculated on structured grids, with values averaged over grid cells that are equidistant from the limbus. To reduce noise caused by the sampling of discrete vessels and tips onto the grids, two Gaussian smoothing passes are applied to *ρ* and *n* before further processing [34].

CPU times for 90 simulated hours on a single processor range from 15 seconds for the Planar2D case, with a fixed VEGF field, to 30 minutes for the Hemisphere and dynamic VEGF model. The most computationally expensive elements of the simulation are the PDE solution times and spatial searches for anastomosis events.

## Results

Fig 4A shows simulated vessel networks after 85 hours (3.5 days) for the case with a fixed VEGF concentration field. Anastomosis is found to be more prevalent in the 2D domains, leading to a reduced number of tips and greater confinement of tips toward the advancing front. In 3D, the presence of multiple vessels through the cornea thickness is evident. In the Circle2D case, there is a tendency for tips to move together as the center is approached, this focusing effect being due to the domain geometry.

**Fig. 4.**
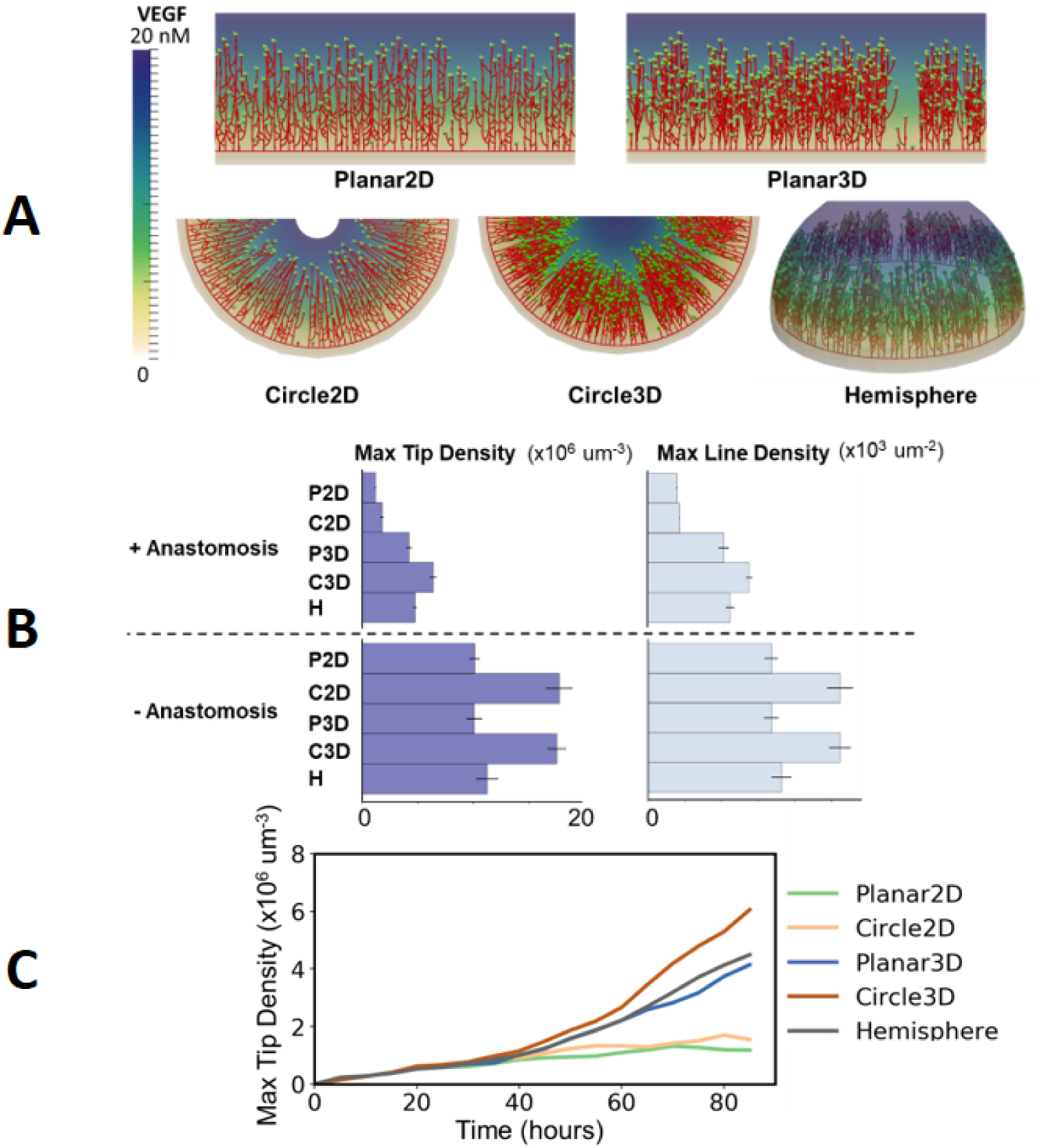
2D and circular domains lead to increased anastomosis. **A)** Simulation results for a fixed VEGF gradient 85 hours after pellet implantation. **B)** A comparison of maximum tip and line density across the studied domains after 85 hours, with and without anastomosis and with *c*_*p*_ = 20 nM. Error bars show one standard deviation from the mean, with five random realisations per observation. **C)** The change in maximum tip density with time in one random realisation per geometry, with anastomosis.

Fig 4B quantifies the maximum tip and vessel line densities in each domain after 85 hours, with and without anastomosis. Without anastomosis, the circular domains have tip densities higher than the planar domains and Hemisphere by a factor of 1.7 due to geometrical effects. Despite the extra vessel length and volume available for sprouting in 3D, line and tip densities are similar to the 2D cases. When anastomosis is active, the tip and line densities in the 2D cases decrease by greater amounts than in 3D. In the planar domain, the tip density decreases by a factor of 5.3 for the Planar2D case but only 1.8 for the Planar3D case. Similarly, it decreases by a factor of 6.4 for the Circle2D case, but only 2.1 for the Circle3D. These results show that the 2D domains lead to greater anastomosis, with the highest tendency for anastomosis in the circular domains. This effect becomes increasingly apparent as the initial pellet concentration *c*_*p*_ is varied from 1 through to 100 nM (shown in S1 Fig). As shown in Fig 4C and the full tip and line density profiles in S2 Fig, the differences between 2D and 3D domains become more pronounced with time, as the capacity for sprouting in 2D is reduced due to anastomosis and lateral inhibition. The increasing density in the circular domains with time is again due to geometric effects.

Fig 5A shows predicted VEGF concentrations in each domain after 1 hour for the dynamic VEGF model. VEGF distributions are quite different across each domain, showing the importance of choosing a suitable representation of the pellet. The Planar2D and Planar3D domains have a relatively high VEGF concentration at the cornea-pellet interface along the entire domain width *W*. When pellets of finite width are used the region of higher concentration is localized to a line of length *w* on the interface. The Planar3D case, with finite pellet width, has a noticeably lower VEGF concentration than the 2D case, due to the pellet thickness *T*_*p*_ being smaller than that of the cornea *T*. In the circular domains, the situation is reversed, with higher VEGF concentrations in the 3D domain due to a greater surface density at the cornea-pellet interface. A higher concentration is observed in the Hemisphere for the same reason.

**Fig. 5.**
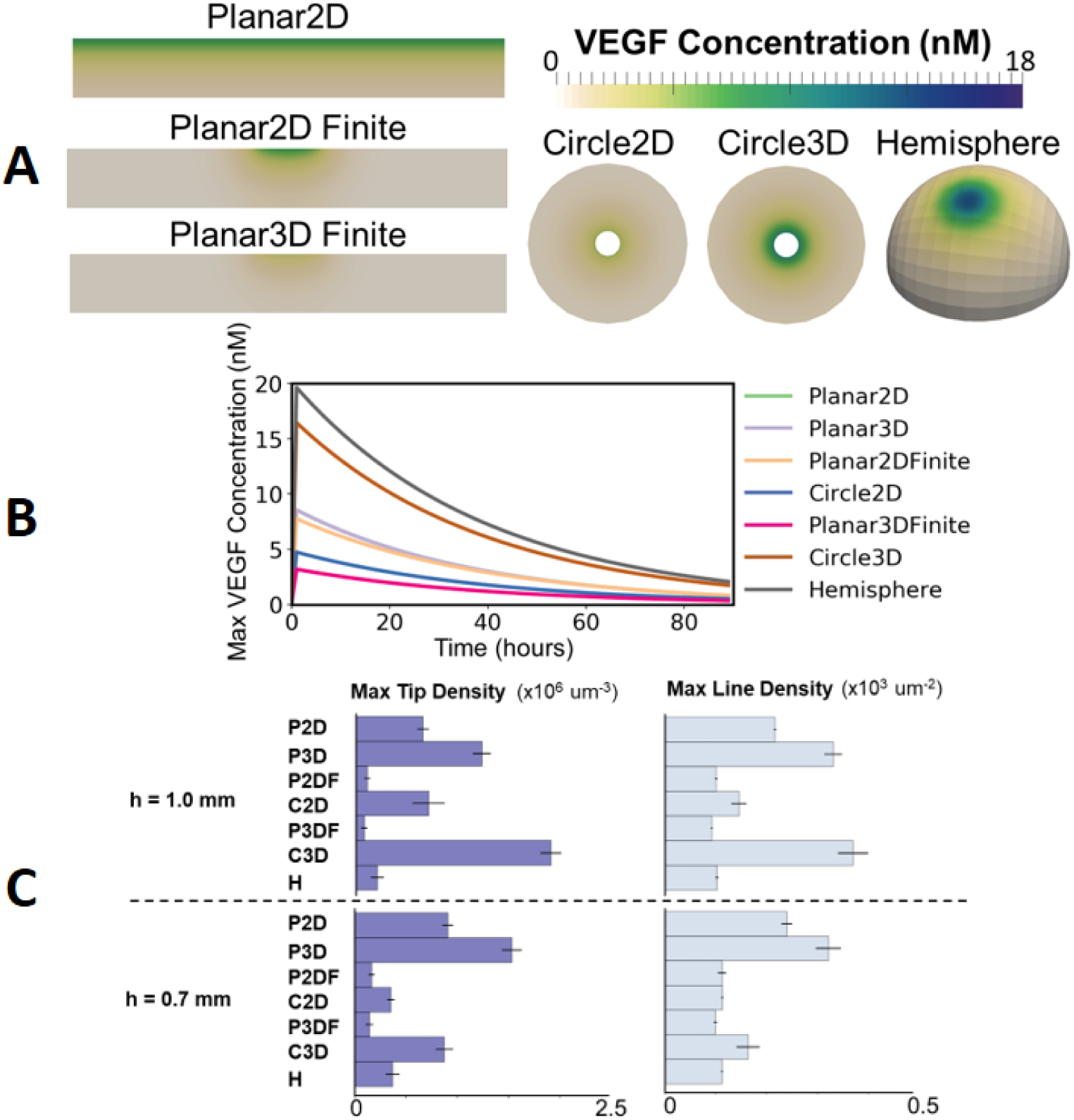
Line and tip density predictions are sensitive to pellet representation. **A)** A comparison of VEGF concentrations for the dynamic model in each domain at 1 hour and *h* = 1.0 mm.**B**) The change in maximum VEGF concentration with time for one set of random realisations of the case with h = 1.0 mm. **C)** A comparison of maximum tip and line densities across the studied domains for *h* = 1.0 mm and *h* = 0.7 mm. Error bars show one standard deviation from the mean, with five random realisations per observation.

Over time the VEGF in the pellet depletes, with the decay term in Eqn (9) being dominant. This leads to a similar rate of decay across all domains, as shown in Fig 5B. The VEGF has largely decayed after 4 days, with a 95 % reduction from the maximum value in the tissue at this time.

The variation in VEGF concentrations shown in Fig 5A, combined with the greater tendency for anastomosis in 2D and circular domains shown in Fig 4, leads to a variety of predicted maximum tip and line densities across the studied domains for the dynamic VEGF model, as shown in Fig 5C. In this case, higher densities are predicted in the planar geometries with extended pellets, while geometries with the finite pellet width have a lower density, more comparable with the Hemisphere, due to focusing of the vascularized region. When the pellet is moved closer to the limbus the general trend is for an increase in the maximum tip density. In relative terms, the tip density increases most in the Hemisphere and Planar3D geometry with finite pellet, both by a factor of 1.7, although in absolute terms the density in the Planar3DFinite geometry is approximately half that of the Hemisphere. In contrast, in the circles, the tip density is reduced by a factor of 1.25 as the pellet is moved towards the limbus. This is a geometric effect, due to the breaking of symmetry as the pellet is moved away from the center of the circle. The dynamics of the maximum densities are similar to those shown in Fig 4C, with the rate of density increase being greatest in the circular domains.

Fig 6 summarizes the locations of the vascular front across all of the studied domains at 85 hours, with activation and de-activation of various biophysical features of the angiogenesis model. The ‘distance of the vascular front to the limbus’, *d* in Fig 1, is found to be insensitive to domain choice, for the dynamic VEGF model shown in Fig 6B, the variation from the Hemisphere value across all domains for the predicted vascular front location is at most 3.7 percent. The lack of sensitivity to domain choice is likely due to: i) the closest tips to the pellet always being near the line of symmetry of all domains, ii) the statistical effect of the metric always accounting for the ‘fastest’ moving vessels amongst the population and iii) the assumption of constant migration speed in the adopted model of tip migration. Increased sensitivity to the domain geometry would be expected if the migration speed depended on VEGF concentration. The location of the maximum tip density and half-maximum tip density are useful additional metrics in cases where they can be measured. As shown in Fig 6C, these metrics are more sensitive to changes in the biophysical mechanisms of network formation than the front location. For example, the tendency for tip cells to be positioned closer to the moving front in 2D domains (shown in Fig 4A) is captured in the bottom glyph in Fig 6C. The effect of strong chemotaxis is similar, as shown in the top glyph. When chemotaxis is relatively weak, or the degree of persistence in the random walk is low, the location of the maximum tip density is moved closer to the limbus. These tendencies are captured across all domains using the maximum tip density and half-maximum tip density location metrics, although they are more sensitive to domain choice than the front location. In all cases the front velocity is approximately constant in time, which is in agreement with experimental observations [7,41], and is similar across all domains.

**Fig. 6.**
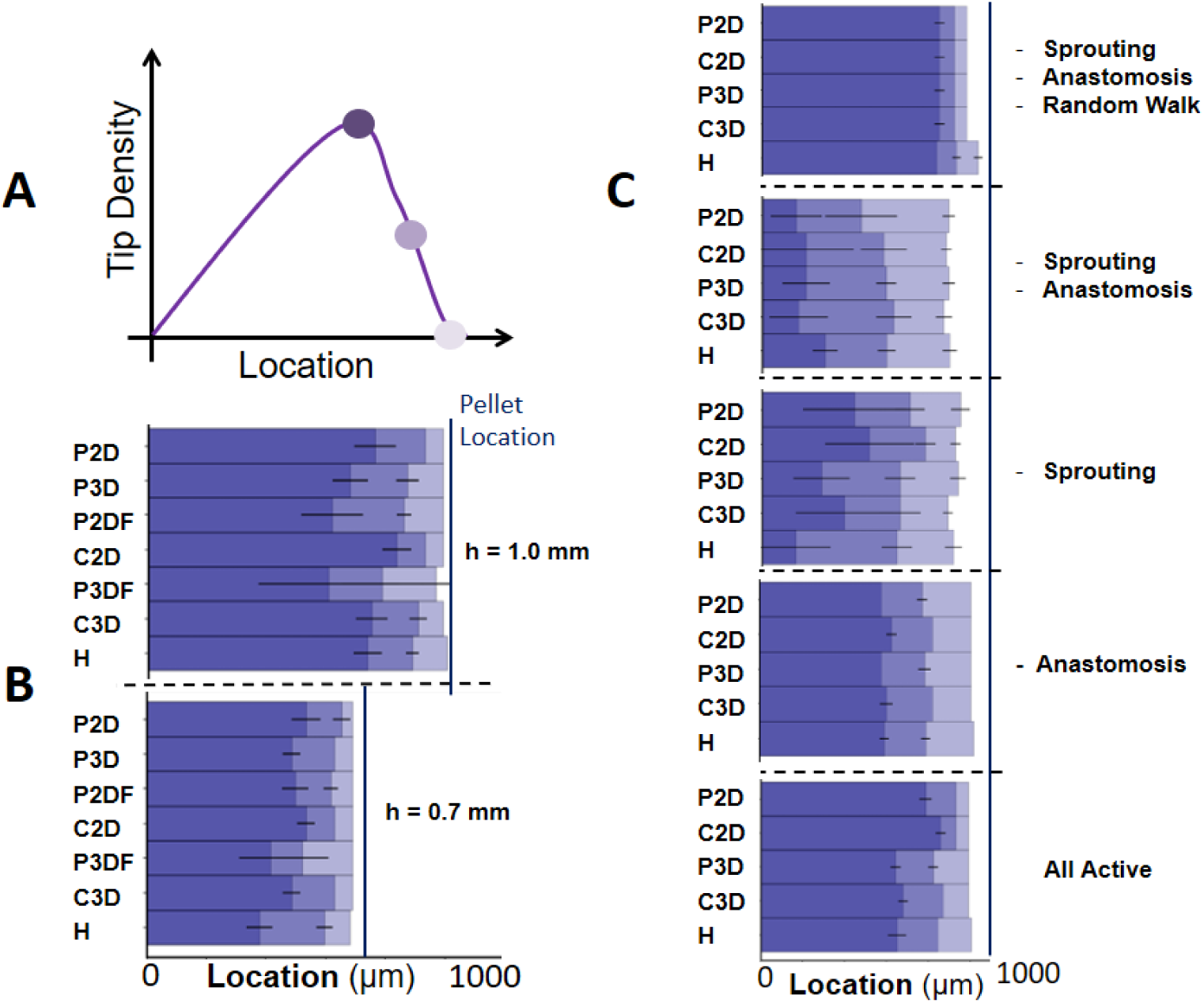
‘Distance to limbus’ is insensitive to domain choice. **A)** A schematic showing measured locations of the tip density profile for comparison in B and C. Locations correspond to the positions of maximum density, half-maximum density and 1% of the maximum density. The ‘distance to limbus’ metric is taken to be equivalent to the location with 1% of the maximum density. **B)** A comparison of tip density profiles for the dynamic VEGF model and different values of *h* at 85 hours. **C)** A comparison of tip density profiles for the fixed VEGF field and activation and de-activation of different biophysical features (indicated by ‘-’). Sprouting is still permitted at the limbus for the ‘- Sprouting’ case. Error bars show one standard deviation from the mean, with five random realisations per observation.

Fig 7A shows the different predicted vessel network patterns in a selection of domains as the pellet is moved closer to the limbus for the dynamic VEGF model. The clear differences in vessel network patterning are not well captured by the ‘distance to limbus’ metric in this case. Although the maximum tip and line density metrics used in Fig 5 are useful in the context of modeling, they can be difficult to measure experimentally. This is because endothelial tip cells are not obvious at typical imaging resolution, while line density measurements are subject to potential errors as multiple vessels may overlap through the cornea thickness [7]. In contrast, it is easier to estimate the ‘vascularized fraction’ or volume of the domain with vessels divided by total domain volume directly from images. In the present study this metric is calculated by accumulating the volume of the cells in the structured grid used in the calculation of densities that are occupied by vessels and dividing by the volume of all cells in the grid.

**Fig. 7.**
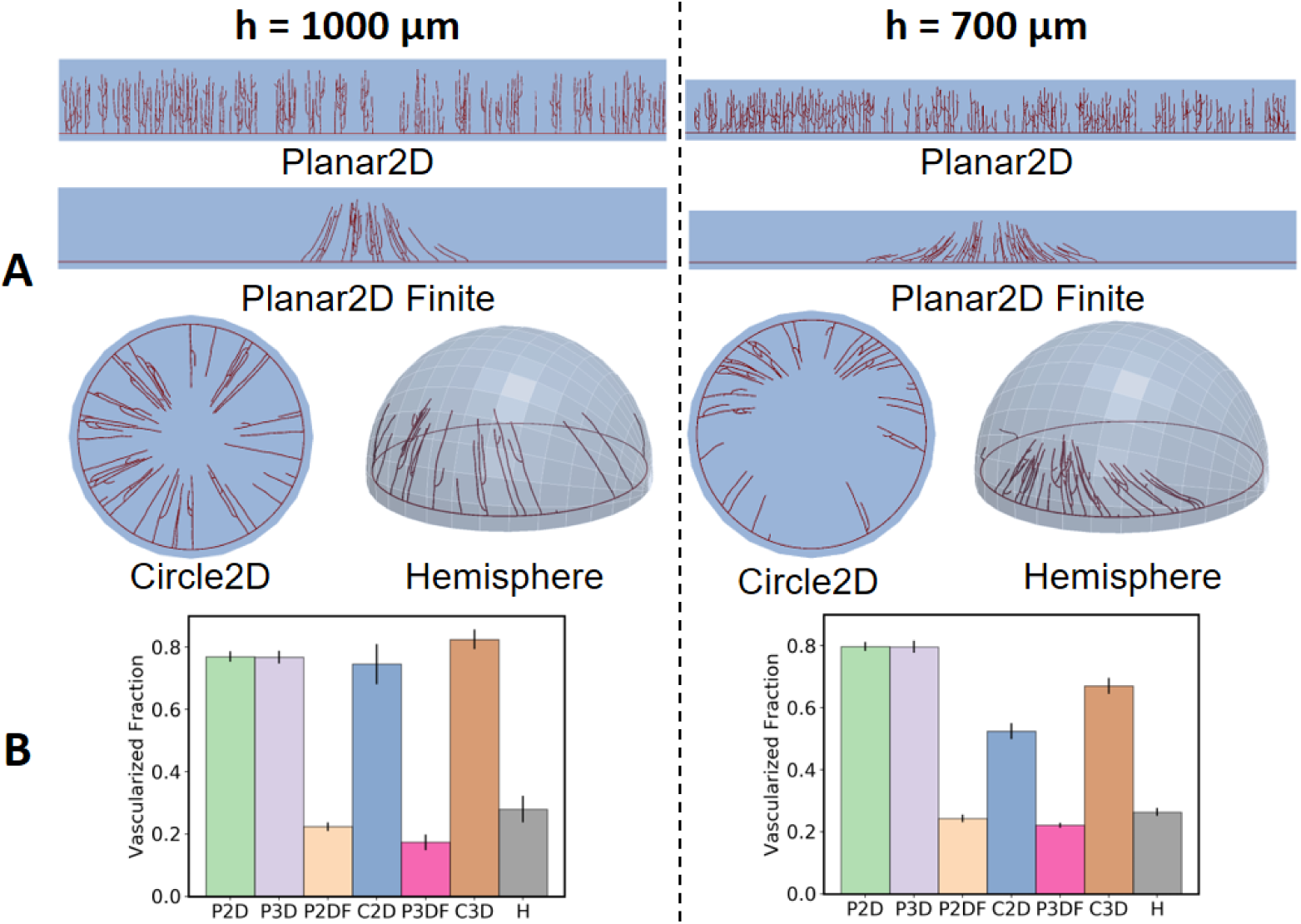
‘Vascularized fraction’ is a useful proxy for tip and line densities. **A)** Simulation results at 85 hours showing different degrees of vascularization in different simulation domains for *h* = 1.0 mm and *h* = 0.7 mm. B) The corresponding ‘Vascularized Fraction’ for each pellet height. Error bars show one standard deviation from the mean, with five random realisations per observation.

As shown in Fig 7B the ‘vascularized fraction’ metric is sensitive to the differences in vascularization between domains shown in Fig 5B, and also captures the trend for increased vascularization when the pellet is moved closer to the limbus in all domains, except for the circles. As such, the metric is predicted to be useful for differentiating neovascularization patterns and translating observations across geometries.

## Discussion

Summarizing the results in Figs 4, 5, 6 and 7, it is predicted that the small, but finite, thickness of the cornea can have an important effect on vessel network formation relative to 2D models, reducing the likelihood of vessels anastomosing and increasing the sensitivity of the predicted vascular response to increases in pellet loading. The representation of the pellet significantly affects neovascularization in all domains, with different degrees of sensitivity to the positioning of the pellet itself also evident across the studied geometries.

Circular domains lead to further increases in anastomosis relative to the hemispherical cornea. This observation has important consequences for studies involving qualitative and quantitative comparisons of vascularization with experiments, such as those mentioned previously [3,6,14,15]. The appearance of ‘brush-border’ effects and the tendency for vessels to approach the pellet, often remarked on in previous studies, occurred naturally in the Circle2D model used in this study, and are largely attributed to the adopted geometry rather than the angiogenesis model. Other occurrences primarily attributed to the circular geometry include the development of artificial loops near the limbus and the grouping of migrating tips toward the vascular front (Fig 4). Direct, quantitative comparisons with experiment and studies on changes in pellet loading or positioning also merit more reflection, given the significant differences in vascularization and sensitivity to pellet positioning between the Circle and Hemisphere geometries shown in Fig 7.

The curvature of the cornea is predicted not to strongly affect neovascularization, with good agreement between the Planar3D domain with a finite pellet and Hemisphere simulations. There is agreement between the 1D models (i.e. Planar2D and Planar3D) and the Hemisphere in terms of predicted front locations, but given the overall insensitivity of this metric to domain choice, it is unclear if a prediction of this quantity alone by the 1D model is particularly informative. For example, the 1D model fails to capture the changing locations of maximum and half-maximum tip densities as the pellet is brought closer to the cornea (see Fig. 6). In fact, it is difficult to interpret how quantities such as vessel line or tip densities should be related between the 1D and 3D models when the vascular front is being focused as per Fig. 7 B. These questions are important in the context of studies such as Connor *et al.* [5], who use 1D models to predict vessel line density profiles for comparison with experiment.

Regarding choice of metrics, it is evident that metrics such as line and tip densities are both difficult to measure and translate across models (as per Fig. 5). The front location (or ‘distance of the vascular front to the limbus’), is straight-forward to measure, but is insensitive to both domain choice and changes to several biophysical mechanisms in the adopted angiogenesis model (Fig. 6). The vascularized fraction is straight-forward to measure and when combined with front location appears to distinguish qualitatively similar vascularization in different domain geometries (Fig. 7).

In addition to giving insights on differences between domain geometries, it is envisaged that the results presented in this study will be useful in the future formulation of PDE models of neovascularization in the cornea. In particular, they give insights into front locations and line and tip density profiles in cases where direct formulation of PDE analogues is challenging, such as accounting for off-lattice random walks with anastomosis and symmetry breaking in the positioning of the spatially resolved pellet. This is particularly the case for the results in Figs 4, S1 Fig and S2 Fig which use a simple, static VEGF profile.

In the present work, a simplified model of neovascularization is adopted, with the primary focus being on comparing predicted neovascularization across different geometries, which has not been performed for the cornea before. By varying phenomenological parameters, such as the chemotactic strength *χ* and deviation in the persistence angle *σ* (as shown in Fig 6), and physical parameters, such as the VEGF concentration in the pellet *c*_*p*_ (as shown in S1 Fig), through a range of values, it is possible to account for a variety of vascular patterning behaviors. Suggested extensions to the adopted modeling are: i) the inclusion of blood flow and vessel regression, which may reduce overall vessel densities and average vessel lengths and ultimately halt the progression of the moving front [42], ii) a more detailed model of endothelial cell proliferation and migration [6 which would remove the need for a phenomenological migration speed *s* and may lead to a front velocity that is not constant in time, iii) models of tip attraction [42] and mechanical interaction with extracellular matrix [6], which may encourage extra anastomosis by guiding tips to locate each other and iv) modeling detailed feedback between a metabolically active tissue and the vasculature [42].

Further, there is scope for more detailed parameter studies, such as an investigation of the relationship between pellet location and ‘vascularized fraction’ and the sensitivity of predictions to the anastomosis radius *r*_*ana*_. More quantitative comparison with experimental measurements of the kind in Tong and Yuan [3] and Connor *et al*. [5] is now possible, due to the focus on modeling the actual geometry of the cornea-pellet system in the present study. Such comparisons would be useful for model validation. In particular, a comparison of experimentally measurable metrics, such as front locations and vascularized fractions, with model predictions while varying pellet loading and positioning would be a useful validation step. Figs 5C and Figs 7B make clear and experimentally testable predictions regarding how the representation of the VEGF pellet affects vessel network formation in different geometries. Exploring these further, via dedicated experiments in a collection of corresponding geometries, would be also be useful.

## Conclusion

In this study we developed a 3D, off-lattice mathematical model to predict neovascularization in a spatially resolved representation of the corneal micropocket assay for the first time. We used the model to study how: i) the geometry of the cornea-pellet system in the micropocket assay affects vessel network formation and ii) which metrics of neovascularization are most sensitive to geometrical differences between typical *in silico*, *in vitro* and *in vivo* tissue domains. We predict that:

- 2D and circular domains lead to increased anastomosis, even relative to the thin cornea geometry, and ultimately different vascular patterning to the spatially resolved model,
- predictions of neovascularization are highly sensitive to the geometrical treatment of the VEGF-containing pellet,
- measuring the distance of the growing vascularized front to the limbus leads to predictions that are insensitive to differences in domain choice, with both positive and negative connotations depending on application,
- vascularized fractions can serve as a useful proxy for densities of migrating tips or vessel line densities, which are more difficult to measure. These metrics can better distinguish underlying vascular patterning than the distance to the limbus alone,
- predictions from planar domains with a finite pellet representation are in closest agreement with those of the hemisphere domain. 3D domains give closer tip and line density predictions to the hemisphere than 2D.

All raw data and software used in this study are available on the Zenodo public archive at doi.org/10.5281/zenodo.995720. Instructions for reproducing the study figures are included in the archive. Source code is available under a BSD-3 Clause license and other data under a Creative Commons CC-BY-4 license.

## Acknowledgements

The research leading to these results has received funding from the European Union’s Seventh Framework Program for Research, Technological Development, and Demonstration under grant No. 600841 (to J.A.G., A.J.C., H.M.B., and J.M.P.-F.). The authors thank Bostjan Markelc and Ruth Muschel of the CRUK/MRC Oxford Institute for Radiation Oncology, University of Oxford for helpful discussions.

## Supporting information

51 Fig. Max tip and line densities as the pellet concentration *c*_*p*_ is increased from 1 to 100 nM.Max tip and line densities after 85 simulated hours for the case with a constant VEGF concentration field. Differences between geometries become more pronounced as the pellet concentration increases.

52 Fig. Full tip and line density profiles sampled every 5 hours for selected domains. Full tip and line density profiles for a single random realisation in a selection of domains for the case with a constant VEGF concentration field.

